# Generalizing RNA velocity to transient cell states through dynamical modeling

**DOI:** 10.1101/820936

**Authors:** Volker Bergen, Marius Lange, Stefan Peidli, F. Alexander Wolf, Fabian J. Theis

## Abstract

The introduction of RNA velocity in single cells has opened up new ways of studying cellular differentiation. The originally proposed framework obtains velocities as the deviation of the observed ratio of spliced and unspliced mRNA from an inferred steady state. Errors in velocity estimates arise if the central assumptions of a common splicing rate and the observation of the full splicing dynamics with steady-state mRNA levels are violated. With scVelo (https://scvelo.org), we address these restrictions by solving the full transcriptional dynamics of splicing kinetics using a likelihood-based dynamical model. This generalizes RNA velocity to a wide variety of systems comprising transient cell states, which are common in development and in response to perturbations. We infer gene-specific rates of transcription, splicing and degradation, and recover the latent time of the underlying cellular processes. This latent time represents the cell’s internal clock and is based only on its transcriptional dynamics. Moreover, scVelo allows us to identify regimes of regulatory changes such as stages of cell fate commitment and, therein, systematically detects putative driver genes. We demonstrate that scVelo enables disentangling heterogeneous subpopulation kinetics with unprecedented resolution in hippocampal dentate gyrus neurogenesis and pancreatic endocrinogenesis. We anticipate that scVelo will greatly facilitate the study of lineage decisions, gene regulation, and pathway activity identification.

## Introduction

Single-cell transcriptomics has enabled the unbiased study of biological processes such as cellular differentiation and lineage choice at single cell resolution ^1,2^. The resulting computational problem is known as trajectory inference. Starting from a population of cells at different stages of a developmental process, trajectory inference algorithms aim to reconstruct the developmental sequence of transcriptional changes leading to potential cell fates. A multitude of such methods have been developed, commonly modeling the dynamics as the progression of cells along an idealized, potentially branching trajectory ^3–8^. A central challenge in trajectory inference is the destructive nature of single-cell RNA-seq, which only reveals static snapshots of cellular states. To move from descriptive towards predictive trajectory models, additional information is required to constrain the space of possible dynamics that could give rise to the same trajectory ^9,10^. As such, lineage-tracing assays can add information via genetic modification to enable the reconstruction of lineage relationships ^11–17^. However, these assays are not straightforward to set up and are technically limited in many systems, such as human tissues.

The concept of RNA velocity has enabled the recovery of directed dynamic information by leveraging the fact that newly transcribed, unspliced pre-mRNAs and mature, spliced mRNAs can be distinguished in common single-cell RNA-seq protocols, the former detectable by the presence of introns ^18^. Assuming a simple per-gene reaction model that relates abundance of unspliced and spliced mRNA, the change in mRNA abundance, termed RNA velocity, can be inferred. The combination of velocities across genes can then be used to estimate the future state of an individual cell. The original model ^18^ estimates velocities under the assumption that the transcriptional phases of induction and repression of gene expression last sufficiently long to reach both an actively transcribing and an inactive silenced steady-state equilibrium. After inferring the ratio of unspliced to spliced mRNA abundance that is in a constant transcriptional steady state, velocities are determined as the deviation of the observed ratio from its steady-state ratio. Inferring the steady-state ratio makes two fundamental assumptions, namely that (i) on the gene level, the full splicing dynamics with transcriptional induction, repression and steady-state mRNA levels are captured; and (ii) on the cellular level, all genes share a common splicing rate. These assumptions are often violated, in particular when a population comprises multiple heterogeneous subpopulations with different kinetics. We refer to this modeling approach as the “steady-state model”.

To resolve the above restrictions, we developed scVelo, a likelihood-based dynamical model that solves the *full* gene-wise transcriptional dynamics. It thereby generalizes RNA velocity estimation to transient systems and systems with heterogeneous subpopulation kinetics. We infer the genespecific reaction rates of transcription, splicing and degradation, and an underlying gene-shared latent time in an efficient expectation-maximization framework. The inferred latent time represents the cell’s internal clock, which accurately describes the cell’s position in the underlying biological process. In contrast to existing similarity-based pseudotime methods, this latent time is grounded only on transcriptional dynamics and accounts for speed and direction of motion.

We demonstrate the capabilities of the dynamical model on various cell lineages in hippocampal dentate gyrus neurogenesis^19^ and pancreatic endocrinogenesis^20^. The dynamical model generally yields more consistent velocity estimates across neighboring cells and accurately identifies transcriptional states as opposed to the steady-state model. It provides fine-grained insights into the cell states of cycling pancreatic endocrine precursor cells, including their lineage commitment, cell-cycle exit, and finally endocrine cell differentiation. Here, our inferred latent time is able to reconstruct the temporal sequence of transcriptomic events and cellular fates. Moreover, scVelo identifies regimes of regulatory changes such as transition states and stages of cell fate commitment. Herein, scVelo identifies putative driver genes of these transcriptional changes. Driver genes display pronounced dynamic behaviour and are systematically detected via their characterization by high likelihoods in the dynamic model. This procedure presents a dynamics-based alternative to the standard differential expression paradigm.

Finally, we propose to further account for stochasticity in gene expression, obtained by treating transcription, splicing and degradation as probabilistic events. We show how this can be achieved for the steady-state model and demonstrate its capability of capturing the directionality inferred from the full dynamical model to a large extent. We illustrate its considerable improvement over the steady-state model while being as efficient in computation time. The dynamical, the stochastic as well as the steady-state model are available within scVelo as a robust and scalable implementation (https://scvelo.org). For the latter two scVelo achieves a ten-fold speedup over the original implementation (velocyto) ^18^.

## Results

### Solving the full gene-wise transcription dynamics at single-cell resolution

As in the original framework ^18^, we model transcriptional dynamics (Fig. 1a) using the basic reaction kinetics described by

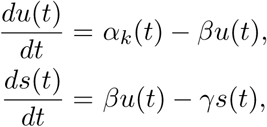

for each gene, independent of all other genes. As opposed to the original framework, to account for non-observed steady states (Fig. 1b), we solve these equations explicitly and infer the splicing kinetics that is governed by two sets of parameters: (i) the reaction rates of transcription *α*_*k*_(*t*), splicing *β*, and degradation *γ*; and (ii) cell-specific latent variables, i.e., a discrete transcriptional state *k*_*i*_ and a continuous time *t*_*i*_, where *i* represents a single observed cell. The parameters of the reaction rates can be obtained if the latent variables are given, and vice versa. Hence, we infer the parameters by expectation-maximization, iteratively estimating the reaction rates and latent variables via maximum likelihood. In the expectation step, for a given model estimate of the unspliced/spliced phase trajectory, 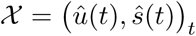, a latent time *t*_*i*_ is assigned to an observed mRNA value *x*_*i*_ = (*u*_*i*_, *s*_*i*_) by minimizing its distance to the phase trajectory 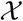 (Fig. 1c). The transcriptional states *k*_*i*_ are then assigned by associating a likelihood to respective segments on the phase trajectory 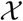, i.e., *k*_*i*_ ∈ {on, off, ss_on_, ss_off_} labeling induction, repression, active and inactive steady states. In the maximization step, the overall likelihood is then optimized by updating the parameters of reaction rates (Fig. 1d, Supp. Fig. 5, Methods).

**Figure 1.**
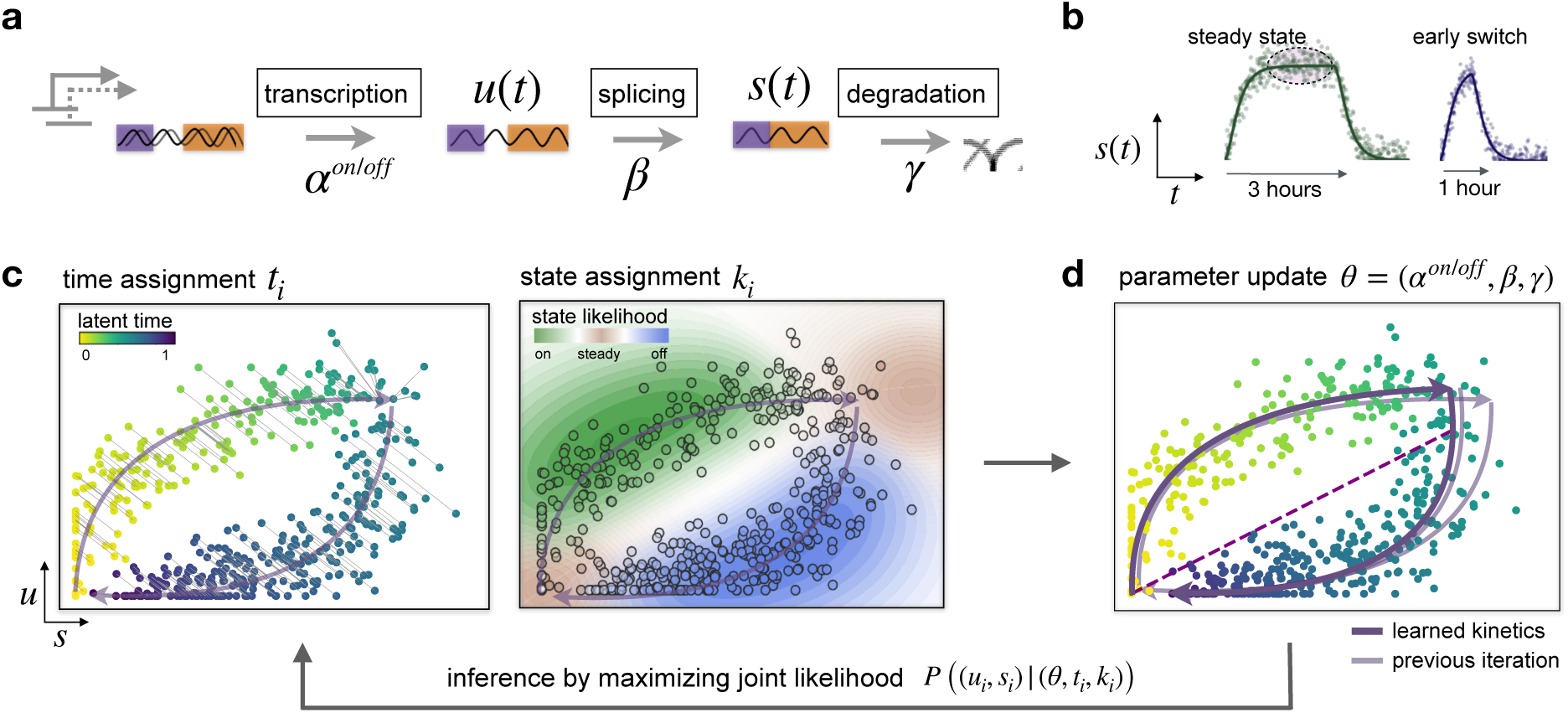
Solving the full splicing kinetics generalizes RNA velocity to transient populations. **a.** Modeling transcriptional dynamics captures transcriptional induction and repression (‘on’ and ‘off’ phase) of unspliced pre-mRNAs, their conversion into mature, spliced mRNAs and their eventual degradation. **b.** An actively transcribed and an inactive silenced steady state is reached when the transcriptional phases of induction and repression last sufficiently long, respectively. In particular in transient cell populations, however, steady states are often not reached as, e.g., induction may terminate before mRNA level saturation, displaying an ‘early switching’ behavior. **c.** We propose scVelo, a likelihood-based model that solves the full gene-wise transcriptional dynamics of splicing kinetics, which is governed by two sets of parameters: (i) reaction rates of transcription, splicing and degradation, and (ii) cell-specific latent variables of transcriptional state and time. The parameters are inferred iteratively via expectation-maximization. For a given estimate of reaction rate parameters, time points are assigned to each cell by minimizing its distance to the current phase trajectory. The transcriptional states are assigned by associating a likelihood to respective segments on the trajectory, i.e. induction, repression, active and inactive steady state. **d.** The overall likelihood is then optimized by updating the model parameters of reaction rates. The dashed purple line links the inferred (unobserved) inactive with the active steady state.

The resulting gene-specific trajectory 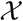, parametrized by interpretable parameters of reaction rates and transcriptional states, explicitly describes how mRNA levels evolve over latent time. While the steady-state model uses linear regression to fit assumed steady states and fails if these are not observed, the dynamical model resolves the full dynamics of unspliced and spliced mRNA abundances and thus enables unobserved steady states to also be faithfully captured (Supp. Fig. 1). RNA velocity is then explicitly given by the derivative of spliced mRNA abundance, parametrized by the inferred variables.

In order to make the inferred parameters of reaction rates relatable across genes, the gene-wise latent times are coupled to a universal, gene-shared latent time that proxies a cell’s internal clock (Suppl. Fig. 2, Methods). This universal time allows us to resolve the cell’s relative position in a biological process with support from the splicing dynamics of all genes. Also transcriptional states can be identified more confidently by sharing information between genes. On simulated splicing kinetics, latent time is able to reconstruct the underlying real time at near perfect correlation and correct scale, clearly outperforming pseudotime. In contrast to pseudotime methods^3,21^, our latent time is grounded on transcriptional dynamics and internally accounts for speed and direction of motion. Hence, scVelo’s latent time yields faithful gene expression time-courses to delineate dynamical processes, and to extract gene cascades.

Further, the coupling to a universal latent time allows us to identify the kinetic rates up to a global gene-shared scale parameter. Employing the overall timescale of the developmental process as prior information, the absolute values of kinetic rates can eventually be identified (Supp. Fig. 3).

### Identifying reaction rates in transient cell populations

To validate the sensitivity of both models with respect to varying parameters in simulated splicing kinetics, we randomly sampled 2,000 log-normally distributed parameters for each reaction rate and time events following the Poisson law. The total time spent in a transcriptional state is varied between two and ten hours.

The ratio inferred by the steady-state model yields a systematic error as the time of transcriptional induction decreases such that mRNA levels are less likely to reach steady-state equilibrium levels (Suppl. Fig. 3a). By contrast, the dynamical model yields a consistently smaller error and is completely insensitive with respect to variability in induction duration. Furthermore, the Pearson correlation between the true and inferred steady-state ratio increases from 0.71 to 0.97 when using the dynamical model. Imposing the overall timescale of the splicing dynamics of 20 hours as prior information, the dynamical model reliably recovers the true parameters of the simulated splicing kinetics, achieving correlations of 0.85 and higher (Supp. Fig. 3b).

### Resolving the heterogeneous population kinetics in dentate gyrus development

To test whether scVelo’s velocity estimates allow identification of more complex population kinetics, we considered a scRNA-seq experiment from the developing mouse dentate gyrus ^19^ (DG) comprising two time points (P12 and P35) measured using droplet-based scRNA-seq (10x Genomics Chromium Single Cell Kit V1, see Methods). The original publication aimed to elucidate the relationship between developmental and adult dentate gyrus neurogenesis. While they successfully linked transient intermediate states to neuroblast stages and mature granule cells, the commitment of radial glia-like cells could not be conclusively determined.

After basic preprocessing, we apply both the steady-state and the dynamical model and display the vector fields using streamline plots^22^ in a UMAP-based embedding ^23^ of the data (Fig. 2a). The dominating structure is the granule cell lineage, in which neuroblasts develop into granule cells. Simultaneously, the remaining population forms distinct cell types that are fully differentiated (e.g. Cajal-Retzius cells) or cell types that form a sublineage (e.g. GABA cells). Whether a cell type is still in transition or already terminal is indicated by two experimental time points and experimental analysis ^19^. Furthermore, the velocities derived from both models settle previously ambiguous evidence of the fate choice of radial glia-like cells in favor of astrocytes over nIPCs.

**Figure 2.**
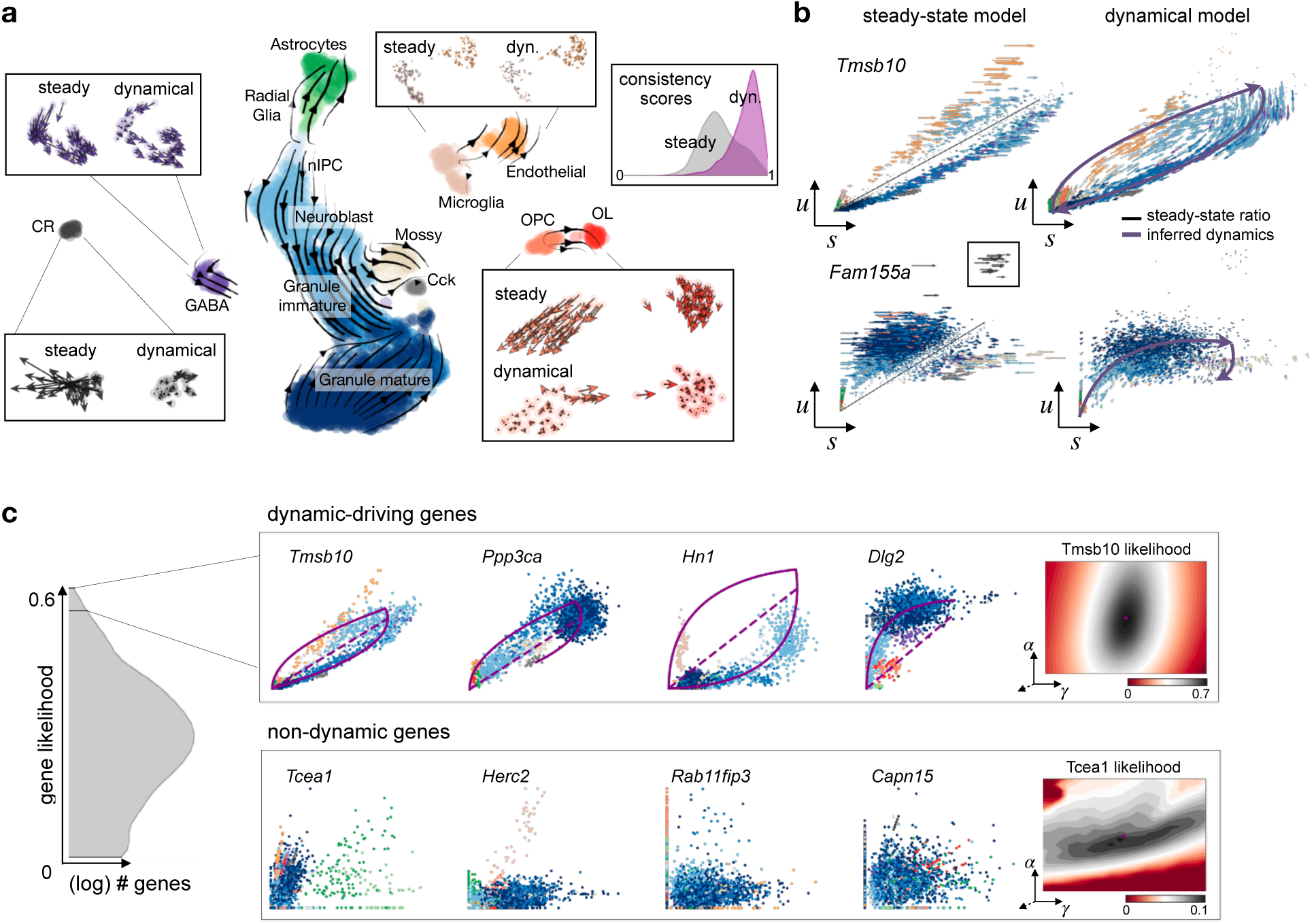
Resolving subpopulation kinetics and identifying dynamical genes in neurogenesis. **a.** Velocities derived from the dynamical model for dentate gyrus neurogenesis^19^ are projected into a UMAP-based embedding. The main gene-averaged flow visualized by velocity streamlines corresponds to the granule lineage, in which neuroblasts develop into granule cells. The remaining populations form distinct cell types that are either differentiated, e.g., Cajal Retzius (CR) cells, or cell types that form sub-lineages, e.g., the GABA and oligodendrocyte lineages (OPC to OL). When zooming into the cell types to examine single-cell velocities, fundamental differences between the velocities derived from the steady-state and dynamical model become apparent. Only the dynamical model identifies CR cells to be terminal by assigning no velocity and indicates that OPCs indeed differentiate into OLs. By contrast, the steady-state model displays a high velocity in CR cells and points OPCs away from OLs. Overall, the dynamical models yields a more coherent velocity vector field as illustrated by the consistency scores (in the top right corner, defined for each cell as the correlation of its velocity with the velocities of neighboring cells). **b.** Gene-resolved velocities allow further interpreting the inferred directionality on the cellular level. For instance, *Tmsb10* is the major contributor to the gene-averaged flow that describes neuroblasts as differentiating into granule cells. With *Fam155a*, the incongruous CR velocities from the steady-state model become evident. By reducing velocity estimation to steady-state deviations, this model is biased to assign high velocities to outlier cells, such as the CR population. In contrast, the dynamical model assign CR cells to a steady state with high likelihoods as they are not well explained by the overall kinetics and cannot be confidently linked to the transient induction state. **c.** The dynamical model allows to systematically identify putative driver genes as genes characterized by high likelihoods. While genes selected by high likelihoods (upper row) display pronounced dynamic behaviour, expression of low-likelihood genes (lower row) is governed by noise or non-existing transient states.

While the main lineage towards mature granule cells is captured by both models, the single-cell velocities illustrate pronounced differences in sub-lineages and sub-clusters. As such, only scVelo correctly identifies the oligodendrocyte precursor cells (OPCs) differentiating into myelinating oligodendrocytes (OLs), and Cajal-Retzius (CR) cells as terminal. The steady-state model erroneously assigns high velocities to CR cells, whic can be traced back to gene-resolved velocities. With *Fam155a*, the incongruous CR velocities from the steady-state model become evident. As the model determines velocities as deviations from steady state that are computed for the whole population, the model is biased to assign high velocities to outlier cells, such as the CR population (Fig. 2b). The dynamical model assign CR cells to steady state with high likelihood as it cannot be confidently linked to any transient state.

*Tmsb10* is the major contributor to the inferred dynamics and illustrates another fundamental difference. Velocities derived from the dynamical model are more consistent across velocities of neighboring cells than those derived from the steady-state model which results in a higher overall coherence of the velocity vector field (Fig. 2a top right, Supp. Fig. 7).

Both the steady-state and the dynamical model yield additional dynamic flow within the mature compartment of granule cells, which was expected to be terminal and may be worthwhile to follow up experimentally.

### Determining dynamical genes beyond differential expression testing

scVelo computes a likelihood for each gene and cell for a model-optimal latent time and transcriptional state, explaining how well a cell is described by the learned spliced/unspliced phase trajectory. Aggregating over cells to obtain an overall gene likelihood, we rank genes according to their goodness-of-fit. This enables us to identify genes that display pronounced dynamic behavior, which makes them candidates for important drivers of the main process in the population (Fig. 2c, Supp. Fig. 4). The top likelihood-ranked genes show clear indication of splicing dynamics, while the expression of low-ranked genes is governed by noise or non-existing transient states. Moreover, partial gene likelihoods, i.e., likelihoods computed for a subset of cells, enable us to identify potential drivers for particular transition phases (induction or repression), branching regions, specific cell types or cycling subpopulations. Many of the top-ranked genes have been reported to play a crucial role in neurogenesis (e.g *Grin2b, Map1b, Dlg2*)^24,25^, while some of these genes were connected to the CA1 region in the hippocampal circuit (e.g. *Tmsb10, Ppp3ca, Hn1*)^26^. By showing that the exclusion of the top likelihood-ranked genes results in non-reconstructability of the dynamics, we also confirm computationally that the inferred directionality is mainly governed by these identified driver genes (Supp. Fig. 6).

### Delineating cycling progenitors, commitment and fate transitions in endocrinogenesis

Next, we demonstrate scVelo’s capabilities to delineate transient lineages in endocrine development in the mouse pancreas, with transcriptome profiles sampled from E15.5^20^. Endocrine cells are derived from endocrine progenitors located in the pancreatic epithelium, marked by transient expression of the transcription factor *Ngn3*. Endocrine commitment terminates in four major fates: glucagon-producing *α*-cells, insulin-producing *β*-cells, somatostatin-producing *δ*-cells and ghrelin-producing *ϵ*-cells ^27^. While in previous works RNA velocity illuminated the directional flow in the endocrine lineage, the endocrine fates could not be clearly delineated, and incongruous subpopulation flows emerged ^20^.

We demonstrate the additional fine-grained insights into the developmental processes that we gain from the dynamical model when compared to the steady-state model. First, scVelo accurately delineates the cycling population of endocrine progenitors (Fig. 3a), biologically affirmed by cell cycle scores((standardized scores of mean expression levels of phase marker genes^28^) and previous analysis ^29^. Further, scVelo illuminates cell states of lineage commitment, cell-cycle exit, and endocrine cell differentiation. By contrast, the steady-state model does not capture the cell cycle and yields incongruous backflows in later endocrine stages (Fig. 3b). For instance, *α*-cells that erroneously seem to be dedifferentiating can be traced back to false state identifications, e.g., in *Cpe* assigning *α*-cells in parts to both induction and repression phase (Fig. 3c).

**Figure 3.**
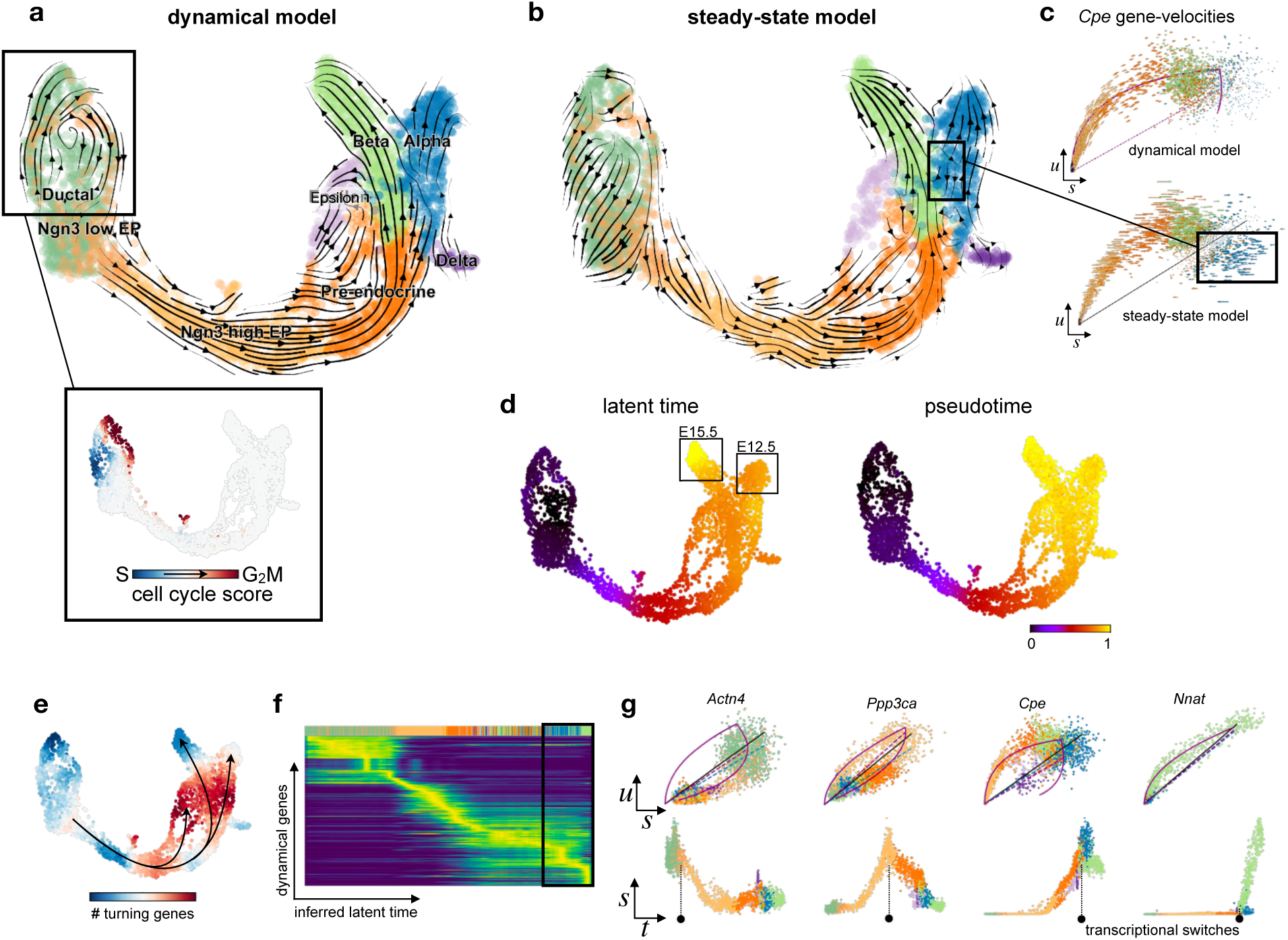
Delineating cycling progenitors, lineage commitment and disentangling cell fates and regimes of transcriptional activity through latent time in pancreatic endocrinogenesis. **a.** Velocities derived from the dynamical model for pancreatic endocrinogenesis^20^ are visualized as streamlines in a UMAP-based embedding. The dynamical model accurately delineates the cycling population of endocrine progenitors (EP), their lineage commitment, cell-cycle exit, and endocrine differentiation. Inferred S and G_2_M phases based on cell-cycle scores affirms the cell cycle identified by the dynamical model. **b.** The steady-state model does not capture the cycle and yields incongruous backflows directed against the lineage in later endocrine stages. **c.** Single-gene velocities illustrate the limitations of the steady-state model. Incongruous backflows in *α*-cells can be traced back to false state identifications, e.g., in *Cpe* assigning *α*-cells in parts to both induction and repression phase. **d.** scVelo’s latent time is based only on transcriptional dynamics and represents the cell’s internal clock. It captures aspects of the actual time better than similarity-based diffusion pseudotime, as observed in the chronology of endocrine cell fates: *α*-cells are produced earlier in actual time (prior to E12.5) while *β*-cells are produced later (E12.5 – E15.5). While latent time enables the temporal relation of the two fates, pseudotime does not distinguish their temporal position. **e.** By using latent time to infer and count switching points between transcriptional states (e.g., from induction to homeostasis), lineage commitment and branching points become apparent. **f.** Gene expression dynamics resolved along latent time shows a clear cascade of transcription in the top 300 likelihood-ranked genes. **g.** Putative driver genes are identified by high likelihoods. Phase portraits (top) and expression dynamics along latent time (bottom) for these driver genes characterize their activity. While *Actn4* switches at cycle-exit and endocrine commitment, the three other genes switch or start to express at the branching points.

### Relating cell fates and disentangling dynamical regimes through latent time

We infer a universal gene-shared latent time that represents the cell’s internal clock. This latent time is a more faithful reconstruction of real time than similarity-based diffusion pseudotime (Supp. Fig. 2, Methods). We compared pseudotime and latent time in the chronology of endocrine cell fates. In real time, *α*-cells are produced earlier (prior to E12.5) than *β*-cells (E12.5-E15.5) ^20^. This ordering is captured by latent time but not by pseudotime (Fig. 3d). Furthermore, the inferred velocities in *α*-cells are lower than the strong directional flow in *β*-cells, which again suggests that *α*-cells have already been produced at an earlier stage. Moreover, the inferred gene-specific switching time points indicate regions of transcriptional changes. The number of identified genes turning from one transcriptional state to another, e.g. from induction to repression, give rise to regions of lineage commitment, transition states and branching points (Fig. 3e). Within these regions, putative driver genes can be identified by their likelihoods, among which many (e.g. *Abcc8, Cpe and Pcsk2*)^30,31^ are known to be associated with insulin regulation. Their transcriptional activity is shown by gene expression dynamics resolved along latent time (Fig. 3f,g).

### Extending the model to account for stochasticity in gene expression

The partial stochasticity of gene expression ^32^ has been addressed by a variety of modeling approaches in systems biology^33^. The flexibility of scVelo’s likelihood-based approach allows us to extend the deterministic ODE model to account for stochasticity by treating transcription, splicing and degradation as probabilistic events. For simplicity, we demonstrate how this can be achieved for the steady-state model (see Methods). The resulting Markov jump process is commonly approximated by moment equations ^34^, which can be solved in closed form in the linear ODE system under consideration. By including second-order moments, we exploit not only the balance of unspliced to spliced mRNA levels but also their covariation. The stochastic steady-state model is capable of capturing the results of the full dynamical model to a greater extent than the deterministic steady-state model (Suppl. Fig. 7a). In particular, the stochastic model displays higher consistency than the deterministic model (Suppl. Fig. 7b), while remaining as efficient in computation time (Suppl. Fig. 9). Investigations of a stochastic dynamical model are left for future work.

### Ten-fold speedup for the steady-state model and large-scale applicability

The dynamical, the stochastic as well as the steady-state model are available within scVelo as a robust and scalable implementation (https://scvelo.org). Illustratively, on pancreas development with 25,919 transcriptome profiles, scVelo runs the full pipeline for the steady-state as well as stochastic model from preprocessing the data to velocity estimation to projecting the data in any embedding in less than a minute (Supp. Fig. 9). That is obtained by memory-efficient, scalable and parallelized pipelines via integration with scanpy ^21^, by leveraging efficient nearest neighbor search ^35^, analytical closed-form solutions, sparse implementation and vectorization. The scVelo pipeline thereby achieves more than a ten-fold speedup over the original implementation (velocyto) ^18^. The full splicing dynamics including kinetic rate parameters, latent time and velocities, is inferred in a longer but practicable runtime of 20 minutes for 1k genes across 35k profiles. As it scales linearly with the number of cells and genes, its runtime is exceeded by velocyto’s quadratic runtime on large cell numbers of 35k and higher. For large cell numbers, also memory efficiency becomes a critical aspect. On an Intel Core i7 CPU with 3.7GHz and 64GB RAM, velocyto cannot process more than 40k cells, while scVelo scales to more than 300k cells.

## Discussion

scVelo enables velocity estimation without assuming either the presence of steady states or a common splicing rate across genes. It maintains the weaker assumptions of constant gene-specific splicing and degradation rates and two transcription rates each for induction and repression. These assumptions might be violated in practice and can be addressed by extending scVelo towards more complex regulations: On the gene level, full-length scRNA-seq protocols, such as Smart-seq2^36^, allow accounting for gene structure, alternative splicing and state-dependent degradation rates. These can be incorporated into scVelo’s likelihood-based inference by adapting the ODE model. In particular, spatial single-cell RNA profiling at transcriptome scale ^37,38^ may provide additional information on relative cell positions necessary to resolve spatial dependencies in gene regulation. Stochastic variability may be leveraged beyond steady state, which has been dubbed as ‘listening to the noise’ and shown to improve parameter identifiability ^39^. Extending the kinetic model to protein translation has been proposed within the steady-state formulation^40^ and can be likewise included into the dynamical model. Metabolic labeling, e.g. using scSLAM-seq ^41,42^, enables the quantification of total RNA levels together with newly transcribed RNA. This additional readout can be easily included into the dynamical model, incorporating varying labeling lengths as additional prior. A further extension would be to couple the single-gene dynamical models to formulate regulatory motifs, which may be inferred by leveraging recent parameter inference techniques for scalable estimation and model selection ^43^. Downstream of scVelo, existing trajectory inference methods may be extended towards informing directionality by robustly integrating velocities to better model cell fate decisions. As such, PAGA ^7^ has made a first suggestion for inferring directed abstracted representations of trajectories through RNA velocity.

Beyond the identification of trajectories and the dynamics of single genes, the dynamic activation of pathways is of central importance. By combining scVelo with enrichment techniques, activated pathways can be inferred in a systematic way, without relying on clustering and differential expression analysis, in analogy to how we demonstrated the inference of dynamically regulated genes. The identification of dynamic pathways and transcription factors immediately lead to testable hypotheses for contributions to cell state transitions. scVelo’s suitability for characterizing transient populations makes it a promising candidate for studying cellular responses to perturbation, which often display drastic switching behaviors. In particular, scVelo could help to mechanistically understand recent machine learning approaches to modeling such response^44^ and point to ways to extend them to incorporate splicing dynamics.

In the meanwhile, scVelo is continuously advanced by the community, bringing efficiency enhancements to the RNA velocity workflow ^45^. It has, for instance, contributed to the detailed study of dynamic processes in human lung regeneration ^46^, and is expected to facilitate the study of lineage decisions and gene regulation, particularly in human.

## Methods

### Preparing the scRNA-seq data for velocity estimation

The raw dataset of hippocampal dentate gyrus neurogenesis^19^ is available in the NCBI Gene Expression Omnibus (GEO) repository, accession number GSE95753. We included samples from two experimental time points, P12 and P35.

The raw dataset of pancreatic endocrinogenesis ^20^ has been deposited under the accession number GSE132188. We included samples from the last experimental time point E15.5.

Annotations of unspliced/spliced reads were obtained using velocyto CLI^18^. Alternatively, reads can be pseudo-aligned with kallisto ^45^.

The datasets are directly accessible in our python implementation (https://scvelo.org).

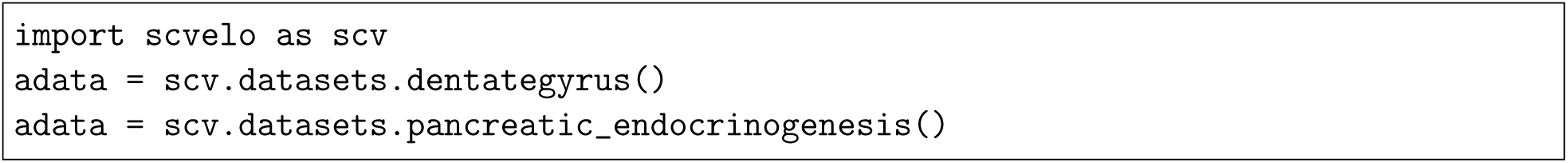

All analyses and results are obtained using default parameters and default data preparation procedures. The count matrices are size-normalized to the median of total molecules across cells. The top 2,000 highly variable genes are selected out of those that pass a minimum threshold of 20 expressed counts commonly for spliced and unspliced mRNA. A nearest neighbor graph (with 30 neighbors) was calculated based on euclidean distances in PCA space (with 30 principal components) on logarithmized spliced counts. For velocity estimation, first and second order moments (means and uncentered variances) are computed for each cell across its 30 nearest neighbors. These are the default procedures in scVelo.

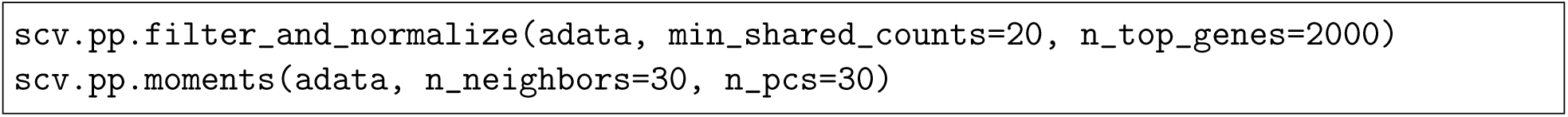

After velocity estimation, the gene space can be further restricted to genes that pass a minimum threshold for the coefficient of determination (*R*^2^, derived from the steady-state model) or gene likelihood (*P* ((*u, s*)*|*(*θ, η*)), derived from the dynamical model).

### Modeling transcriptional dynamics

On the basis of the dynamical model of transcription shown in Fig. 1, we developed a computational framework for robust and scalable inference of RNA velocity. In the following, we first briefly outline the problem of modeling splicing kinetics, explain the steady-state model and, thereafter, describe the novel dynamical model.

The model of transcriptional dynamics captures **transcriptional induction** (“on” phase) and **repression** (“off” phase) of unspliced precursor mRNAs *u*(*t*) with state-dependent rates *α*^(*k*)^, its **splicing** into mature mRNAs *s*(*t*) with rate *β* (i.e. removing introns from pre-mRNAs and joining adjacent exons to produce spliced mRNAs) and eventual **degradation** with rate *γ*, i.e.

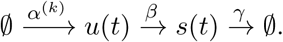

Assuming splicing and degradation rates to be constant (time-independent), we obtain the gene-specific rate equations

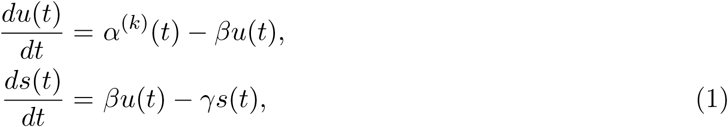

which describe how the mRNA abundances evolve over time. The time derivative of mature spliced mRNA, termed RNA velocity, is denoted as 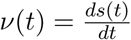.

The quantities *u*(*t*) and *s*(*t*) are size-normalized abundances of unspliced and spliced mRNA, respectively, for a cell measured at time point *t*. In general, the sampled population is not time-resolved and *t* is a latent variable. Likewise, the cell’s transcriptional state *k* is a latent variable that is not known, and the rates *α*^(*k*)^, *β* and *γ* are usually not experimentally measured.

### Steady-state model

Under the assumption that we observe both transcriptional phases of induction and repression, and that these phases last sufficiently long to reach a transcribing (active) and a silenced (inactive) steady-state equilibrium, velocity estimation can be simplified as follows: In steady states, we obtain on average a constant transcriptional state where 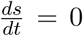 which, by solving Eq. 1, yields 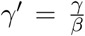 as the steady-state ratio of unspliced to spliced mRNA. It indicates where mRNA synthesis and degradation are in balance. Steady states are expected at the lower and upper quantiles in phase space, i.e. where mRNA levels reach minimum and maximum expression, respectively. Hence, the ratio can be approximated by a linear regression on these extreme quantiles.

It can be solved analytically via a least square fit and is given by

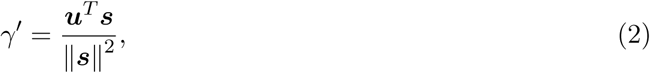

where ***u*** = (*u*_1_, …, *u*_*n*_) and ***s*** = (*s*_1_, …, *s*_*n*_) are vectors of size-normalized unspliced and spliced counts for a particular gene, that lie in the lower or upper extreme quantile, i.e. *n* is only a fraction of the total number of cells. A positive offset can be included into the least square fit to account for basal transcription. The steady-state ratio is then given by 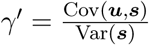, and the offset is given by 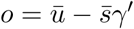, where 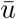 and 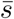 are the means of ***u*** and ***s***, respectively.

Then, velocities can be computed as deviations from this steady-state ratio, i.e.

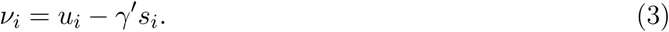

While a constant transcriptional state is reflected by zero velocity, the direction and relative speed during a dynamic process is given by the sign and magnitude of non-zero velocity.

Taken together, under this simplified model, velocities are estimated along two simple equations as steady-state deviations. With this notion, the cumbersome problem of estimating latent time is circumvented. Further, velocities only depend on one ratio instead of absolute values of kinetic rates, which technically corresponds to measuring all entities in units of splicing rate, thus effectively assuming one common splicing rate *β* = 1 across all genes.

scVelo hosts an efficient estimation procedure of velocities derived from the steady-state model.

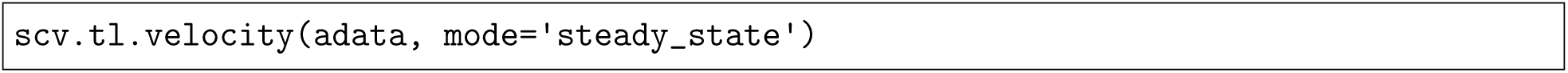

### Dynamical model

#### Model description

In recognition that steady states are not always captured and that splicing rates differ between genes, we establish a framework that does not rely on these restrictions. The analytical solution to the gene-specific rate equations in Eq. 1 is found by integration, which yields

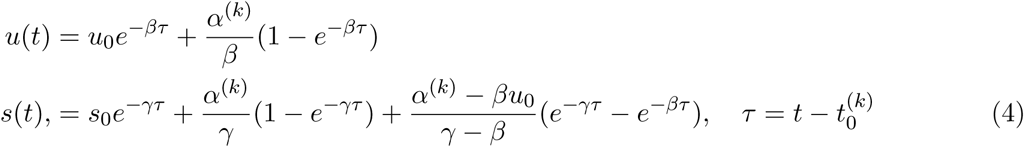

with parameters of reaction rates *θ* = (*α*^(*k*)^, *β*, *γ*), cell-specific time points *t* ∈ (*t*_1_, …, *t*_*N*_), and initial conditions *u*_0_ = *u*(*t*_0_), *s*_0_ = *s*(*t*_0_).

Gene activity is orchestrated by transcriptional regulation, implying that gene up- or down-regulation is inscribed by alterations in the state-dependent transcription rate *α*^(*k*)^. That is, *α*^(*k*)^ can have multiple configurations each encoding one transcriptional state. For the model, this requires an additional parameter set, assigning a transcriptional state to each cell, i.e. *k* = (*k*_1_, …, *k*_*N*_). Consequently, not only *α*^(*k*)^ but also the initial conditions 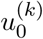, 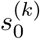 are state-dependent, as well as the time point of switching states 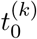. Consider a transition from one state *k* = *i* to a subsequent state *k* = *i* + 1, e.g. from induction to homeostasis. Then, the initial conditions of the next state are given by evaluating the trajectory of the current state at its respective switching time point

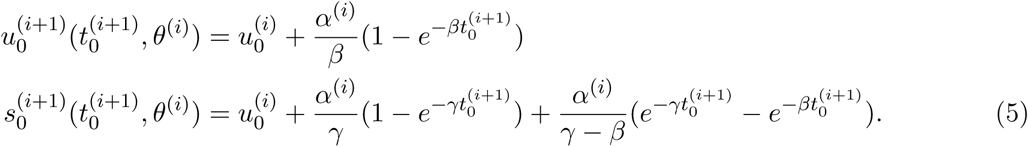

Being at state *k*=*i*, abundances can potentially reach their steady state in the limit,

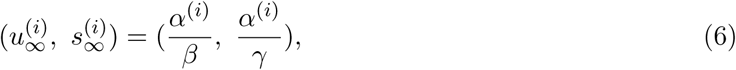

where the number of potential steady states equals the number of transcriptional states.

#### Parameter inference

In the following, we consider four phases, induction (*k*=1), and repression (*k*=0) each with an associated potential steady state (*k*=*ss*_1_, *k*=*ss*_0_). Recovering the splicing kinetics entails inferring the model parameters, i.e. reaction rates *θ*^(*k*)^, time point *t*_*i*_ for each cell that couples the measurement to the system of differential equations by assignment onto the phase trajectory, state *k*_*i*_ to which each cell is assigned and switching time point 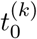 of transitioning to another state.

Let the model estimate be 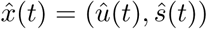, and let the observations be 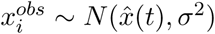. With the assumption of the gene-specific *σ* to be constant across cells within one transcriptional state, and the observations to be i.i.d., the likelihood-based framework is derived in the following.

The negative log-likelihood to be minimized is given by

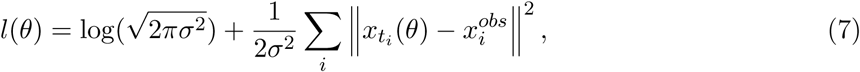

where *θ* = (*α*^(*k*)^, *β*, *γ*).

The cell-specific latent time points are required for coupling an observation to the system of differential equations in order to obtain a mapping of 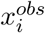 to 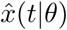. Hence, solving for kinetic rates relies on also estimating latent time which illustrates the problem complexity. This is solved by expectation-maximization (EM) iterating between finding the optimal parameters of kinetic rates and the latent variables of time and state, initialized with parameters derived from the steady-state model, i.e. *β* = 1 and 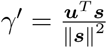, where ***u*** and ***s*** are size-normalized unspliced and spliced counts from extreme quantile cells. The cell-specific state is initialized to be induction or repression depending on whether the sample lies above or below the steady-state ratio, respectively, i.e.

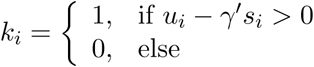

The transcription rates are initialized with 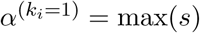 and 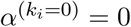.

After initializing the system with meaningful parameters, the expectation-maximization (EM) algorithm iteratively applies the following two steps:

- E-step: Given 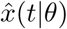 parametrized by the current estimate of *θ*, we assign a latent time *t*_*i*_ to the observed value 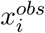 by minimizing the distance to the phase trajectory 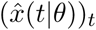 in each transcriptional state. State likelihoods are then assigned to each cell, which yield an expected value of the log-likelihood (by integrating over all possible outcomes for transcriptional states).
- M-step: The parameter set *θ* is updated to maximize the log-likelihood.

We explicitly model both transient states of induction and repression as well as (unobserved) active and inactive steady states. The state likelihoods are determined by the distance of the observations to the four segments of the phase trajectory, parametrized by kinetic rates and latent time.

For latent time, we adopt an explicit formula that approximates the optimal time assignment for each cell. This is applied throughout the EM framework mainly for computation efficiency reasons, while (exact) optimal time assignment is applied in the last iterations. The approximation of optimal latent time is obtained as follows.

The equation for spliced mRNA levels can be rewritten as a function of unspliced mRNA levels:

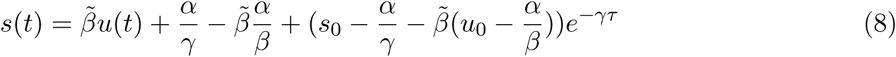

where 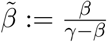 constitutes the linear dependence of unspliced on spliced mRNA.

If we denote 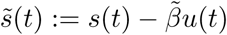 and 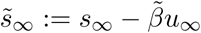, the equation can be rewritten as

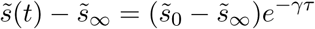

where *τ* can be solved explicitly for each cell by taking the inverse

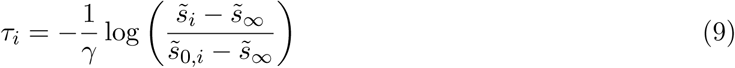

If *β > γ*, the time assignment is thus obtained as inverse of a positive linear combination of unspliced and spliced mRNA dynamics. For genes with *β ≤ γ*, we can instead directly take the inverse of *u*(*t*) which is given by

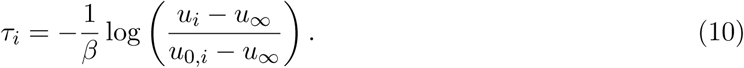

This explicit time assignment is used throughout the parameter fitting, while in the last iteration latent time is solved optimally likelihood-based.

scVelo provides a flexible and efficient module to estimate reaction rates and latent variables.

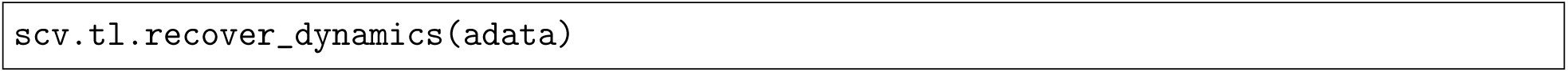

RNA velocity can then be estimated using the explicit description of inferred splicing kinetics.

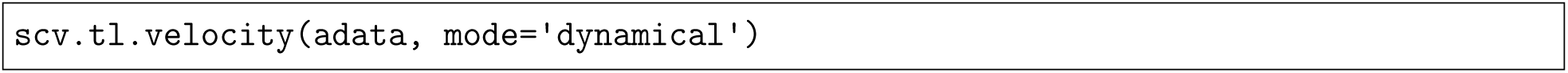

#### Computing transition probabilities from velocities

Assuming that velocities truthfully describe the actual dynamics locally, we estimate transition probabilities of cell-to-cell transitions. Let ***S*** ∈ ℝ ^*n×d*^ be the gene expression matrix of *d* genes across *n* cells. Further, we have estimated the velocity vectors (***ν***_*i*_)_*i*=1,…,*n*_ in the previous section, of which ***ν***_*i*_ ∈ ℝ^*d*^ predicts the change in gene expression of cell ***s***_*i*_ ∈ ℝ^*d*^.

Cell ***s***_*i*_ is expected to have a high probability of transitioning towards cell ***s***_*j*_ when the corresponding change in gene expression ***δ***_*ij*_ = ***s***_*j*_ − ***s***_*i*_ matches the predicted change according to the velocity vector ***v***_*i*_. We apply cosine similarity, i.e. cosine of the angle between two vectors,

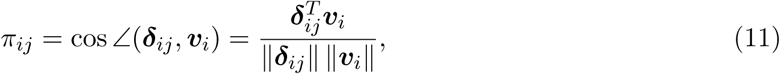

where *π*_*ii*_ = 0. It solely measures similarity in directionality, not in magnitude, and ranges from −1 (opposite direction) over 0 (orthogonal, thus maximally dissimilar) to 1 (identical direction). The resulting similarity matrix ***π*** encodes a graph, which we refer to as *velocity graph*. Optionally, a variance stabilizing transformation can be applied such that the cosine correlation is computed between 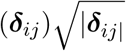 and 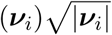.

An exponential kernel is applied in order to transform the cosine correlations into transition probabilities

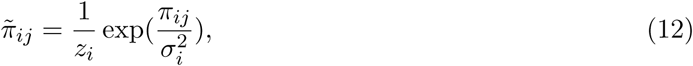

with row normalization factors 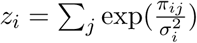 and kernel width parameters *σ*_*i*_ optionally adjusted for each cell locally (across neighboring cells).

The transition probabilities are aggregated into a *transition matrix* 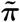 describing the Markov chain of the differentiation process. Throughout the literature, 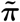 is also referred to as transport map which serves as a coupling of a developmental process. A distribution of cells ***µ*** = (*µ*_1_, …, *µ*_*n*_) can be pushed through the transport map to obtain its descendant distribution. Reversely, a distribution ***µ*** can be pulled back through the transport map to obtain its ancestors, i.e.

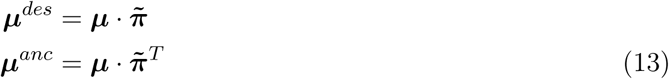

A descendant or ancestor distribution of a set of cells 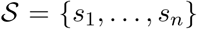 can be obtained by setting 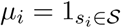, where 1 denotes the indicator function.

scVelo efficiently computes the velocity graph by sparse and vectorized implementation.

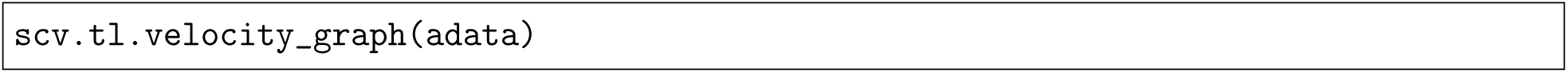

#### Gene-shared latent time

After inferring parameters of kinetic rates, a gene-shared latent time is computed as follows. First, gene-specific time points of well-fitted genes (with a likelihood of at least 0.1) are normalized to a common overall timescale. The root cells of the differentiation process are obtained by computing the stationary states ***µ***^∗^ satisfying

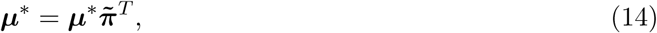

which is simply given by the left eigenvectors of 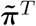 corresponding to an eigenvalue of 1.

Now, for every root cell *o*, we compute the *p*-quantile 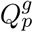 of all respective gene-specific time increments across all genes *g*

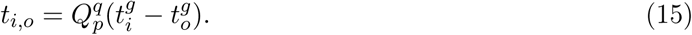

where *p* is chosen such that it maximizes the correlation between the resulting gene-shared latent time-course *t*_0,*o*_,…, *t*_*N,o*_ and its convolution across local neighborhood of cells. The rationale behind taking the *p*-quantile is the adaption to a non-uniform density of cells along the time-course as cells often tend to accumulate in later time points. We find optimal values for *p*, lower than the median, at around 20 − 30%. A high correlation with the convolution of latent time improves robustness and consistency in the estimate.

That is, for each root cell we find the respective time increments that achieve best overall accordance with the learned dynamics and yield local coherence.

Gene-shared latent time of cell *i* is then obtained as mean across all root cells

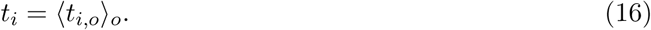

Finally, for robustness, by regressing the gene-shared latent time-course against its neighborhood convolution, we detect inconsistent time points and replace them with their convolutions.

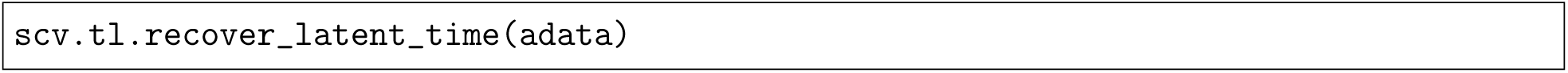

#### Projection of velocities into the embedding

The projection of velocities into a lower-dimensional embedding (e.g. UMAP ^23^) for a cell *i* is obtained on the basis of a transition matrix 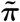(see previous section) which contains probabilities of cell-to-cell transitions that are in accordance with the corresponding velocity vectors,

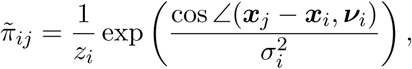

with row normalization factors 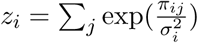 and kernel width parameters *σ*_*i*_.

The positions of cells in an embedding, such as t-SNE or UMAP, are described by a set of vectors 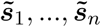. Given the normalized differences of the embedding positions 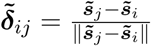, the embedded velocity is estimated as the expected displacements w.r.t. the transition matrix

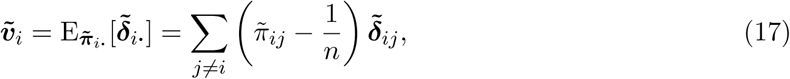

where subtracting 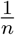 corrects for the non-uniform density of points in the embedding.

The directional flow is visualized as single-cell velocities or streamlines in any embedding.

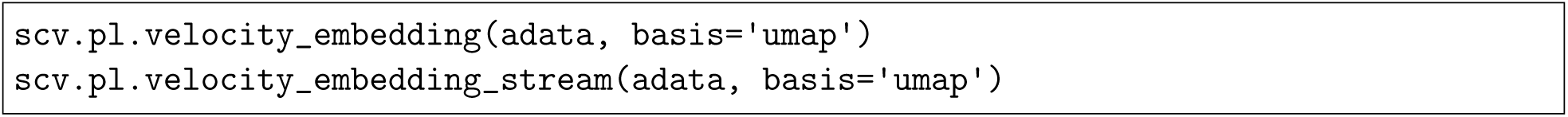

#### Accounting for stochasticity through second-order moments

The model for velocity estimation can be extended with higher order moments, obtained by treating transcription, splicing and degradation as probabilistic events. In this regard, the probabilities of all possible reactions corresponding to these events occurring within an infinitesimal time interval (*t, t* + *dt*] are provided as follows:

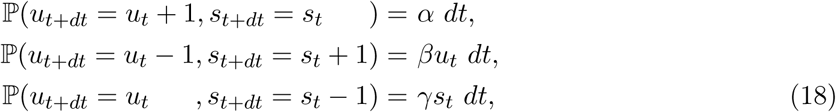

where we denoted *u*_*t*_ = *u*(*t*), *s*_*t*_ = *s*(*t*) to facilitate clarity.

From (18) the time derivative for the uncentered moment 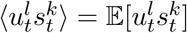 is derived as

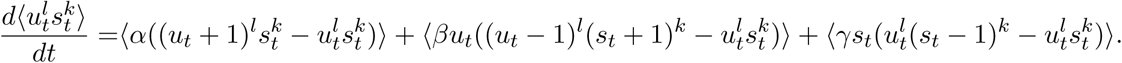

Hence, the firstand second-order dynamics are given by

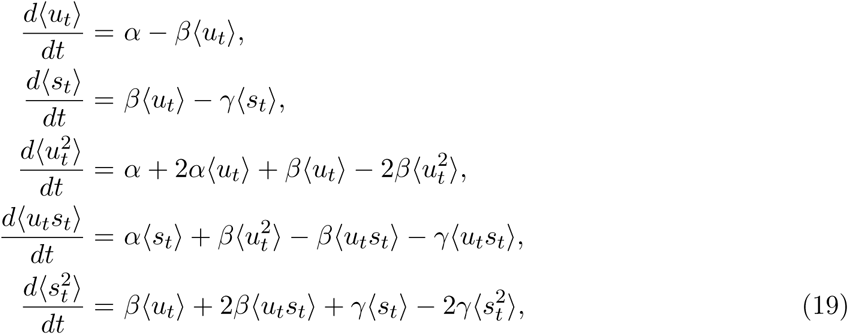

The moments for each cell are computed among a preset number of nearest neighbors of the corresponding cell.

This extensions can be easily applied to the steady-state model. Using both first- and second-oder moments, the steady-state ratio is obtained from the system

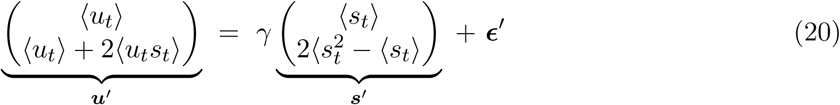

where *E*[***ϵ′***|***s′***] = 0 and Cov[***ϵ′***|***s′***] = *Ω*.

The steady-state ratio can be solved explicitly by generalized least squares and is given by

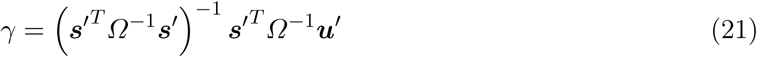

The stochastic model thereby exploits not only the relationship between unspliced and spliced mRNA abundances, but also their covariation.

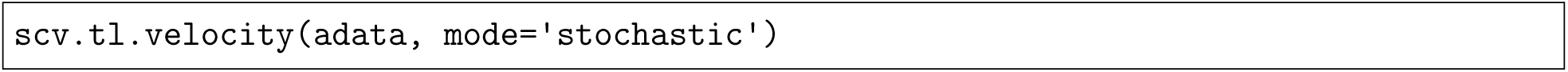

#### Validation metrics

To validate the coherence of the velocity vector field, we define a consistency score for each cell *i* as the mean correlation of its velocity *ν*_*i*_ with velocities from neighboring cells,

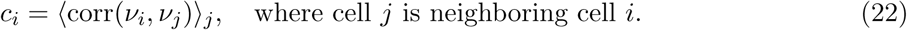

To validate the contribution of a selection of genes (e.g. top likelihood-ranked genes) to the overall inferred dynamics, we define a reconstructability score as follows: The velocity graph consisting of correlations between velocities and cell-to-cell transitions (see previous sections), is computed once (i) including all genes yielding ***π***, and once (ii) only including the selection of genes yielding ***π′***. The reconstructability score is defined as the median correlation of outgoing transitions from cell *i* to all cells that it can potentially transition to, i.e.,

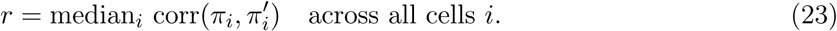

## Code availability

The paper results and our python implementation are available at https://scvelo.org.

## Acknowledgements

We thank P. Kharchenko and S. Linnarsson for stimulating discussions, M. Luecken for valuable feedback on the manuscript and S. Tritschler for valuable feedback on the biological applications. This work was supported by the BMBF grants (01IS18036A and 01IS18053A), by the German Research Foundation (DFG) within the Collaborative Research Centre 1243, Subproject A17, by the Helmholtz Association (sparse2big and ZT-I-0007) and by the Chan Zuckerberg Initiative DAF (182835). M. Lange further acknowledges financial support by the DFG through the Graduate School of QBM (GSC 1006), by the Joachim Herz Stiftung and by the Bayer Foundations.

## Contributions

VB designed and developed the method, implemented scVelo and analyzed the data. FJT conceived the study with contributions from VB and FAW. VB, FAW and FJT wrote the manuscript with contributions from the coauthors. SP contributed to developing scVelo, and ML contributed to developing validation metrics. All authors read and approved the final manuscript.

## Ethics declaration

The authors declare no competing interests.

## Supplementary

**Supllementary Figure 1.**
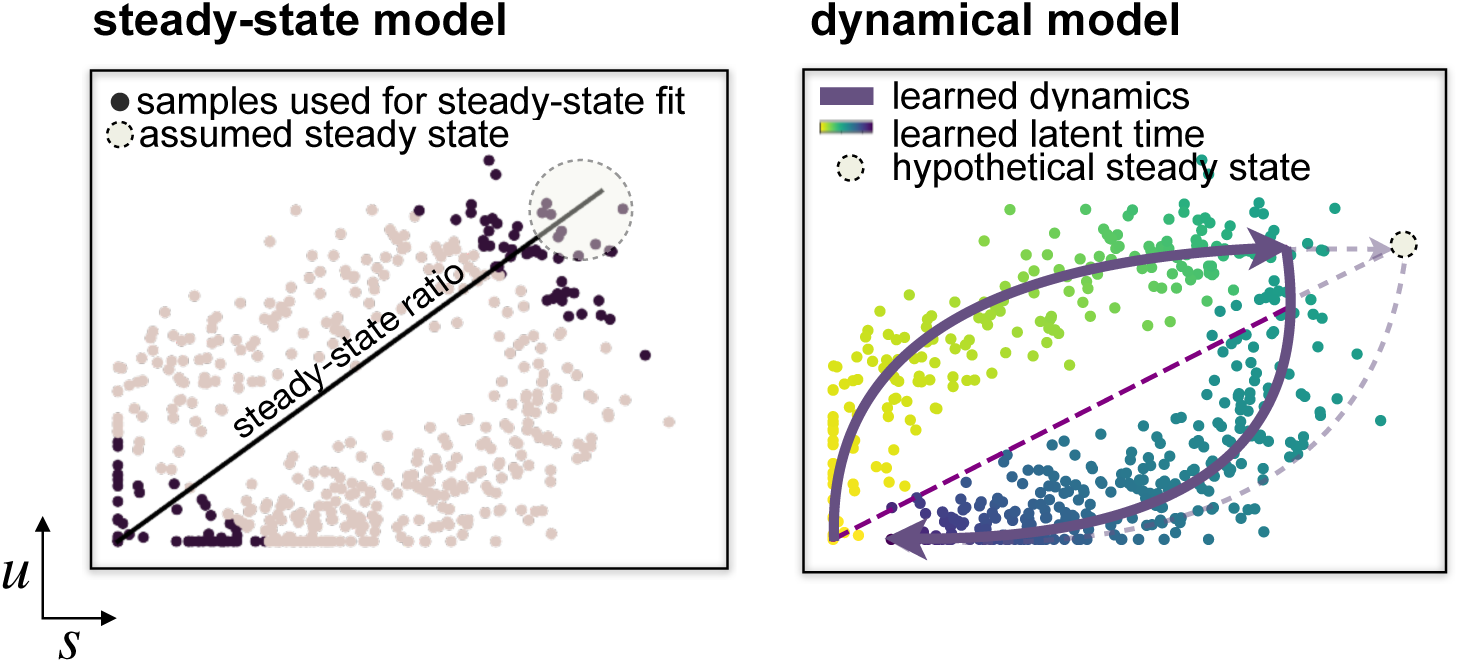
Model fit comparison. The steady-state model determines velocities by quantifying how observations deviate from a ratio of unspliced to spliced mRNA describing an assumed steady-state equilibrium. This steady-state ratio is obtained by performing a linear regression restricting the input data to the extreme quantiles. By contrast, the dynamical model solves the full splicing kinetics and thereby captures non-observed steady states. The error made by the steady-state model for such cases becomes evident when comparing the slope of the line describing the steady-state ratio.

**Supllementary Figure 2.**
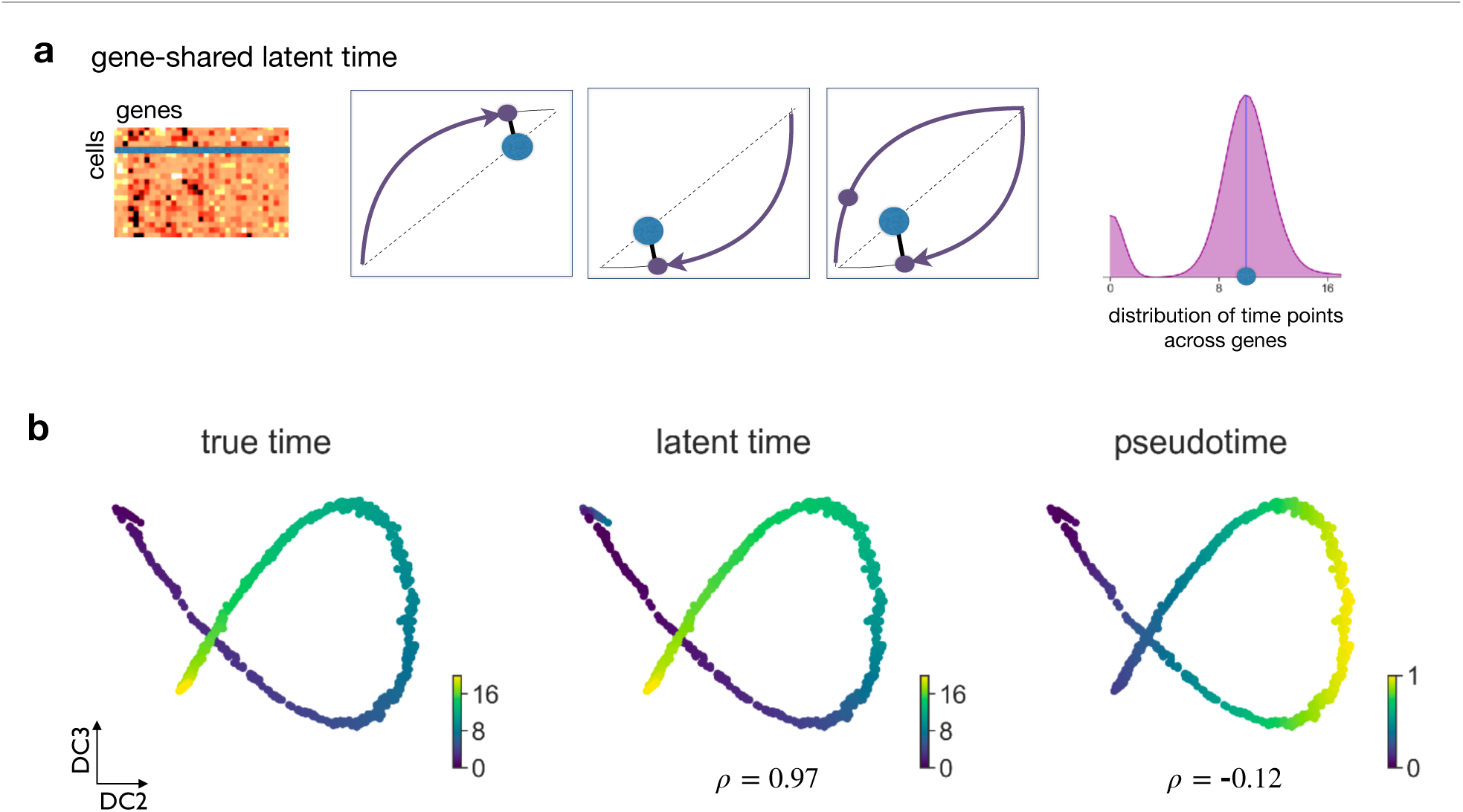
Inferring a universal gene-shared latent time. **a.** A shared latent time is constructed by coupling gene-wise latent times to an additional one-dimensional time covariate across genes. For instance, if a cell is confidently assigned to a later time point in two genes (left) while being ambiguously positioned in a third gene (right), the decision for the later time point is imposed by the majority. Thereby, the cell’s position in the third genes can be confidently identified by sharing information across genes. **b.** The latent time recovered by the dynamical model faithfully captures the underlying real time. While latent time yields a near-perfect Pearson correlation, pseudotime is unable to reconstruct real time.

**Supllementary Figure 3.**
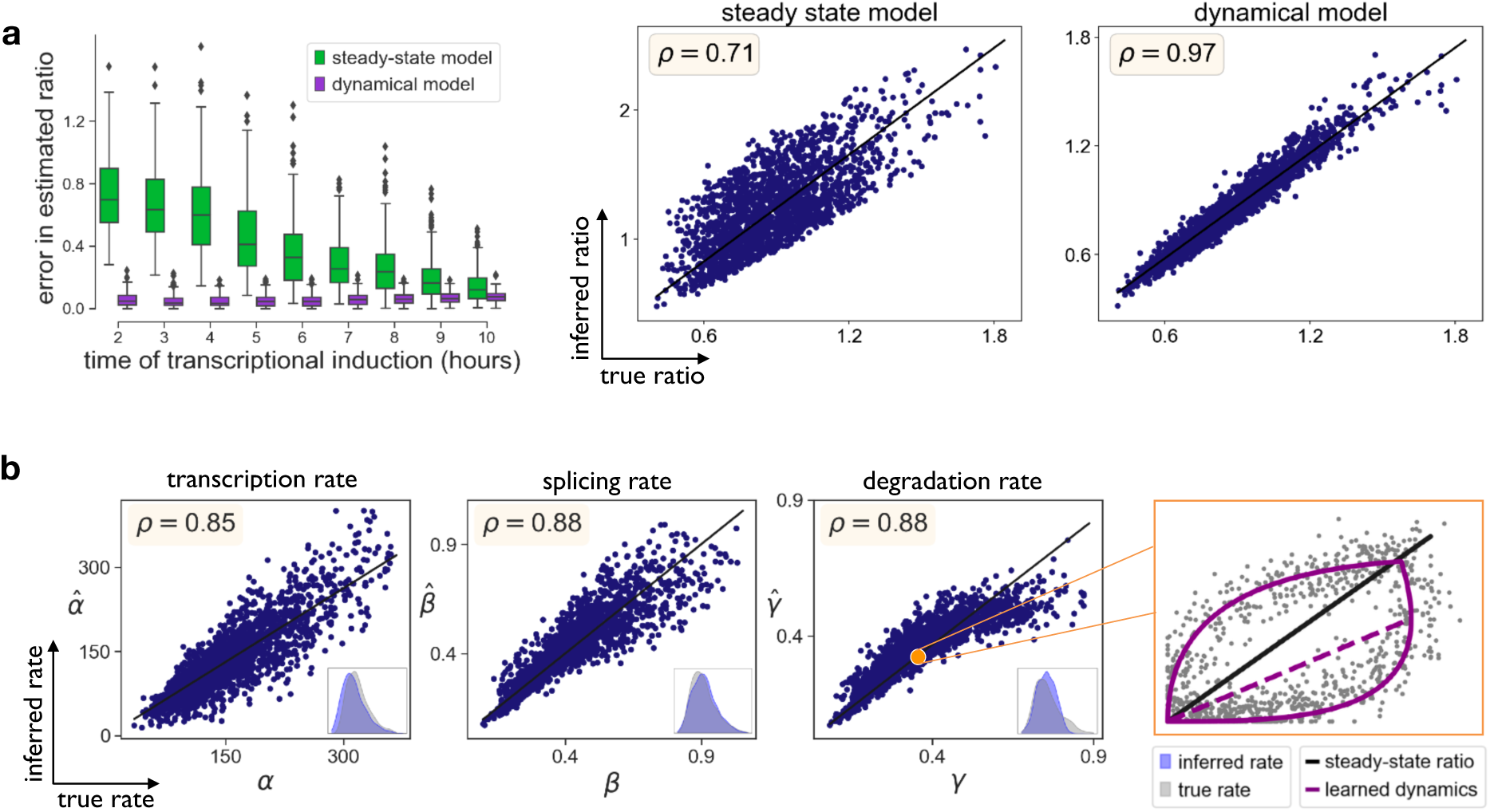
Identifying parameters of reaction rates in transient cell populations. **a.** To validate the sensitivity of both models with respect to varying parameters in simulated splicing kinetics, we randomly sampled 2,000 log-normally distributed parameters for each reaction rate and time events following the Poisson law. The time point of the transcriptional switch (here: from induction to repression) was varied between 2 and 10 hours. As transcriptional induction progresses the abundances converge to an equilibrium steady-state. However, for shorter duration where no equilibrium state has yet been reached, the steady-state model systematically fails to captures the true steady-state ratios. In contrast, the likelihood-based dynamical model consistently infers the true ratio regardless of induction time. The Pearson correlation between the true and inferred steady-state ratio increases from 0.71 to 0.97 when using the dynamical model. **b.** The dynamical model reliably recovers the true parameters of the splicing kinetics. For all kinetic rates (transcription, splicing and degradation), we obtain a correlation of at least 0.85 across the range of 2,000 parameter values for each rate. By imposing an overall timescale of 20 hours as prior information, the dynamical model recovers the absolute values of the reaction rates.

**Supllementary Figure 4.**
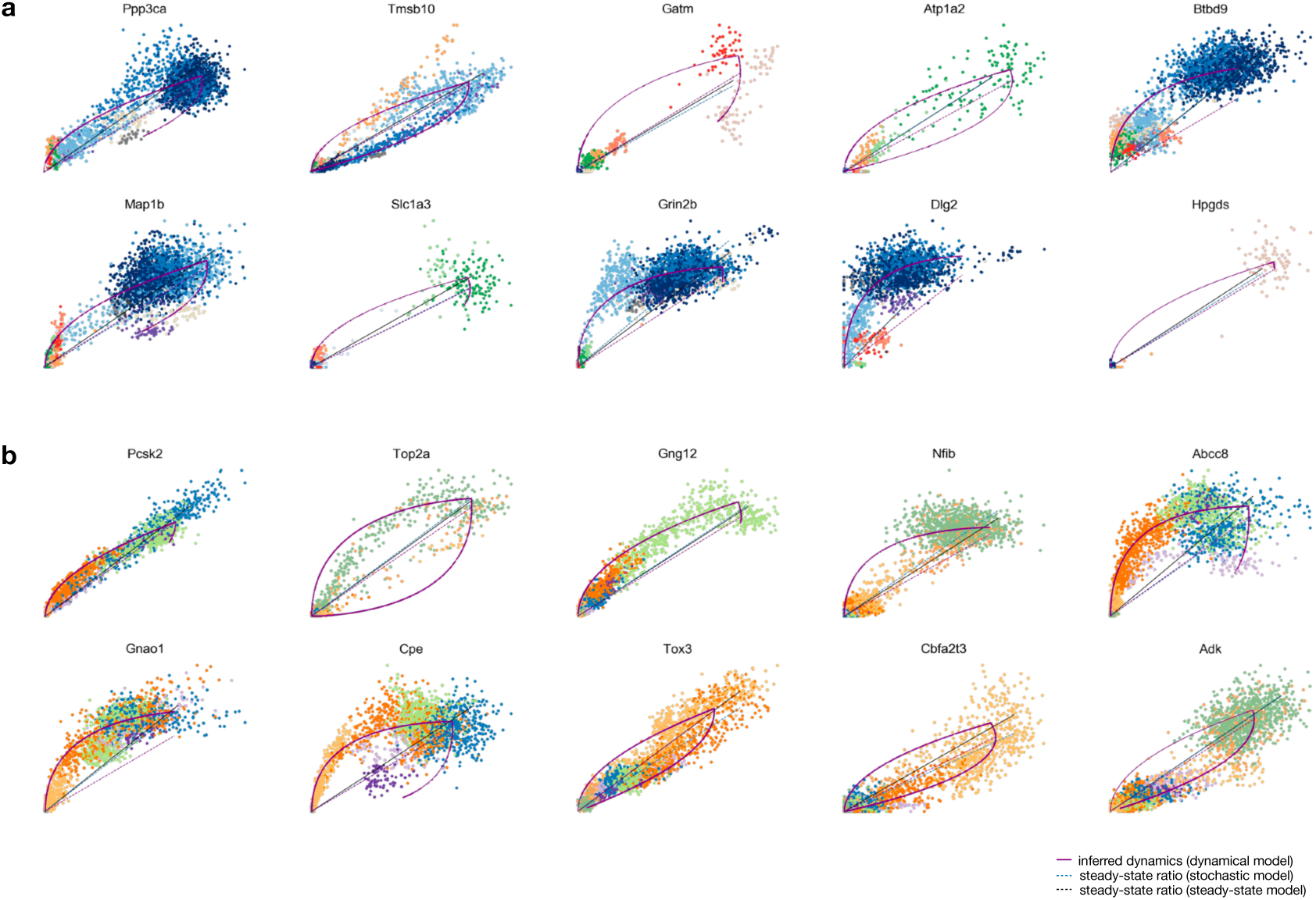
Identified driver genes with pronounced dynamic behaviour. **a.** Top likelihood-ranked genes for hippocampal neurogenesis. **b.** Top likelihood-ranked genes for pancreatic endocrinogenesis.

**Supllementary Figure 5.**
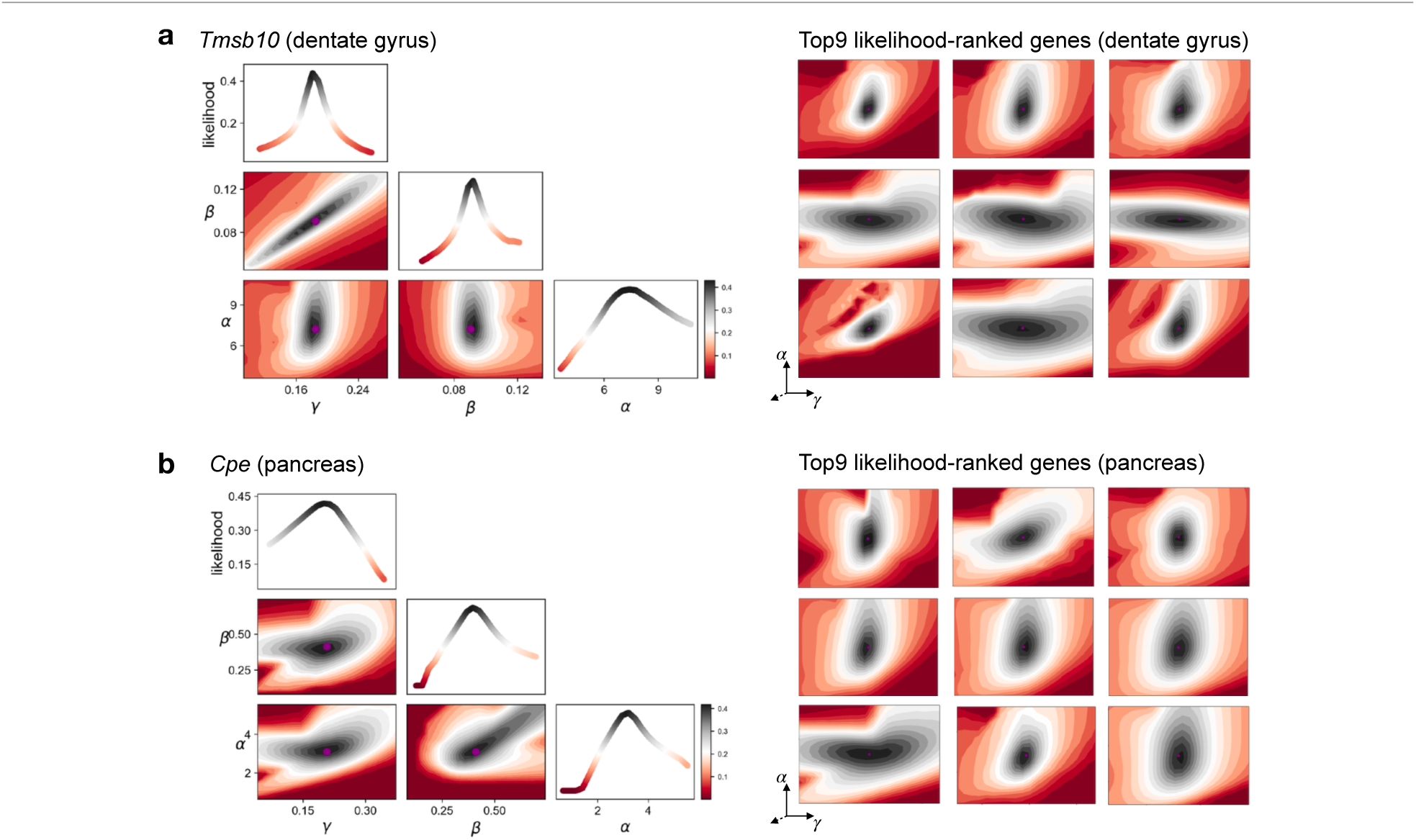
Convergence illustrated by log-likelihood contours. **a.** Objective log-likelihood contours (the optimal value is black colored) for dentate gyrus neurogenesis. Left: Combination of all reaction rates of transcription *α*, splicing *β* and degradation *γ*. Right: transcription and splicing rate for top likelihood-ranked genes. **b.** Objective log-likelihood contours for pancreatic endocrinogenesis.

**Supllementary Figure 6.**
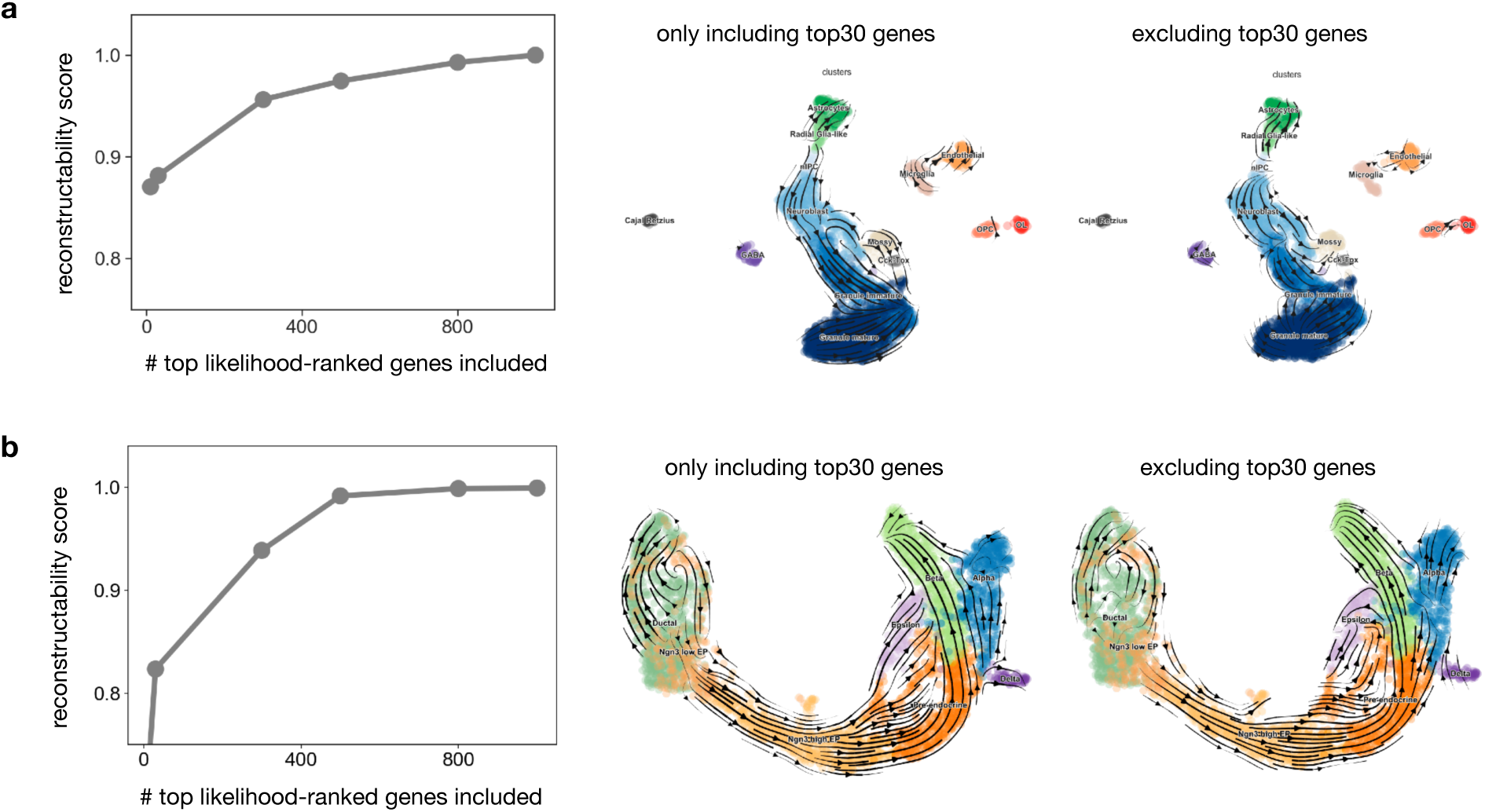
Computational validation of top likelihood-ranked genes. **a.** To validate the contribution of the top likelihood-ranked genes to the overall inferred dynamics, we defined a reconstructability score: it is given by the median correlation of the velocity graph that results from (i) including all genes and (ii) only including a subset of likelihood-ranked genes (see Methods for details). For dentate gyrus, the top 30 genes almost fully explain the inferred dynamics. If we include all other genes and only exclude the top 30 genes, the dynamics cannot be reconstructed. This confirms that these systematically identified genes are transcriptionally highly relevant, making them candidates for important drivers of the main process in the population. **b.** In the pancreatic endocrinogenesis dataset, the reconstructability score similarly confirms that the top likelihood-ranked genes are sufficient to explain the dynamics. However, as opposed to the dentate gyrus example, a much larger set of genes explains the dynamics. This is indicated by the reconstructability curve that starts lower. It is further affirmed as the dynamics that can still be reconstructed when excluding the top 30 genes.

**Supllementary Figure 7.**
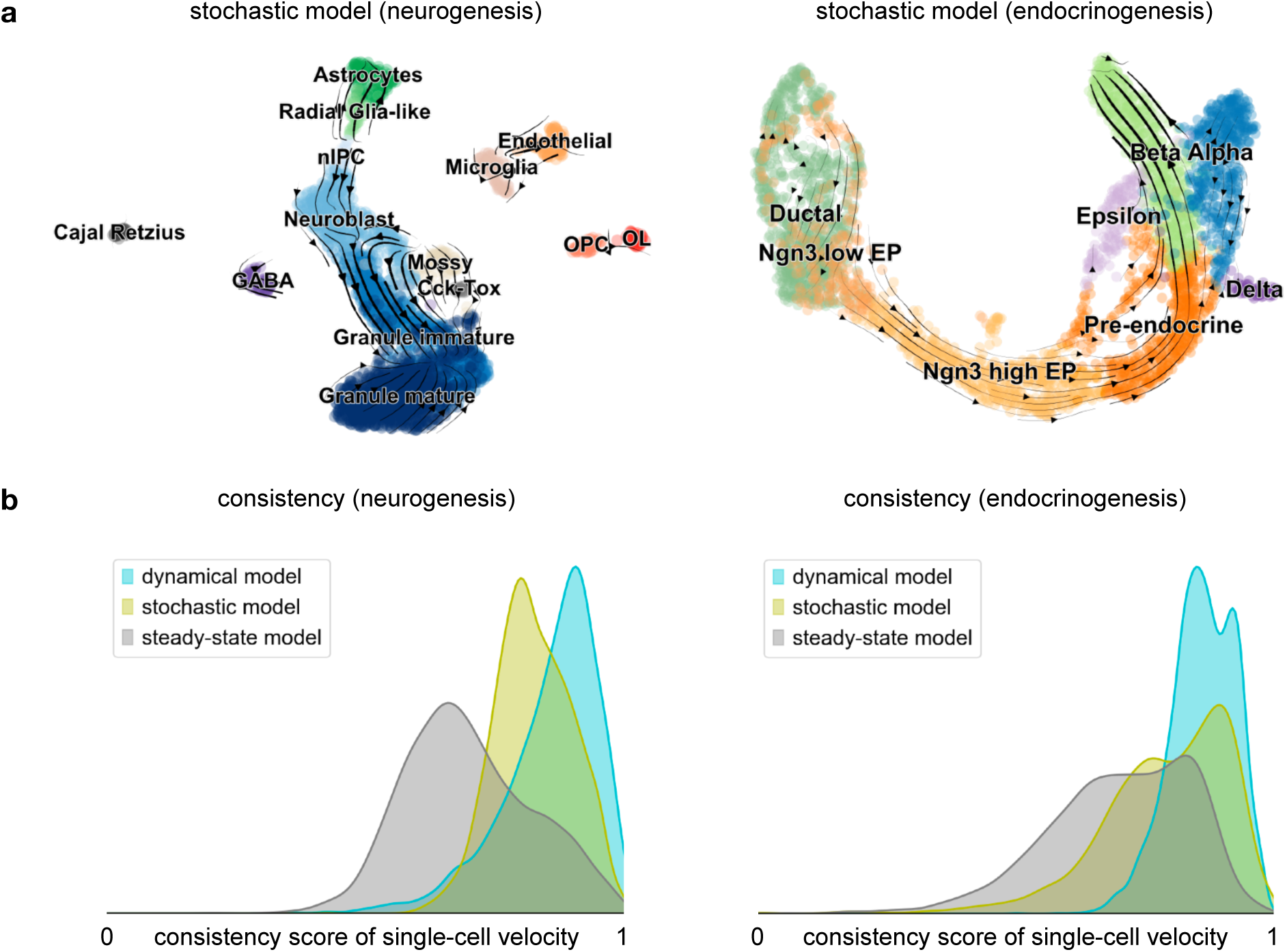
Consistency of single-cell velocities inferred by the stochastic model. **a.** The stochastic model captures the dynamics inferred by the dynamical model to a large extent for dentate gyrus neurogenesis (left) and pancreatic endocrinogenesis (right). It shows great resemblance in dentate gyrus, indicates cycling endocrine progenitors and resolves the endocrine lineage. However, like the steady-state model, the stochastic model also induces a backflow in the *α*-cells. **b.** The consistency score is defined for each cell as the correlation of its velocity with the velocities of neighboring cells. As expected, both in dentate gyrus and pancreas, the stochastic model yields consistencies of single-cell velocities higher than for the steady-state model and lower than for the dynamical model.

**Supllementary Figure 8.**
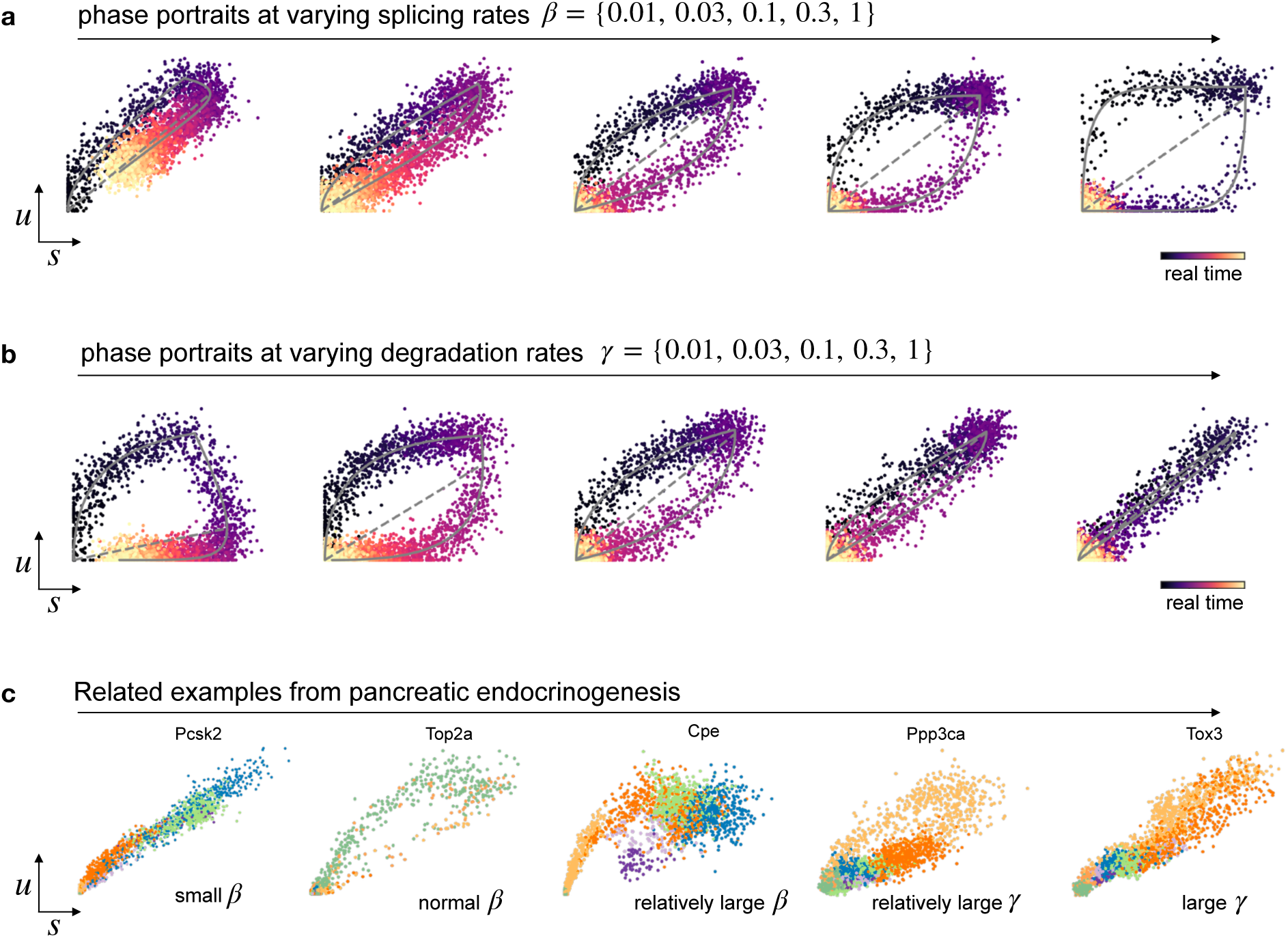
Splicing kinetics at varying reaction rates. **a.** Gene unspliced/spliced phase portraits for simulated kinetics at varying splicing rates show that the curvature increases and the kinetics becomes more pronounced with increasing splicing rate. **b.** The curvature increases with decreasing degradation rates. **c.** Phase portraits of selected top likelihood-ranked genes in pancreatic endocrinogenesis show how the genes from real data mirror the simulated kinetics, which can be explained by the inferred parameters of splicing and degradation rates.

**Supllementary Figure 9.**
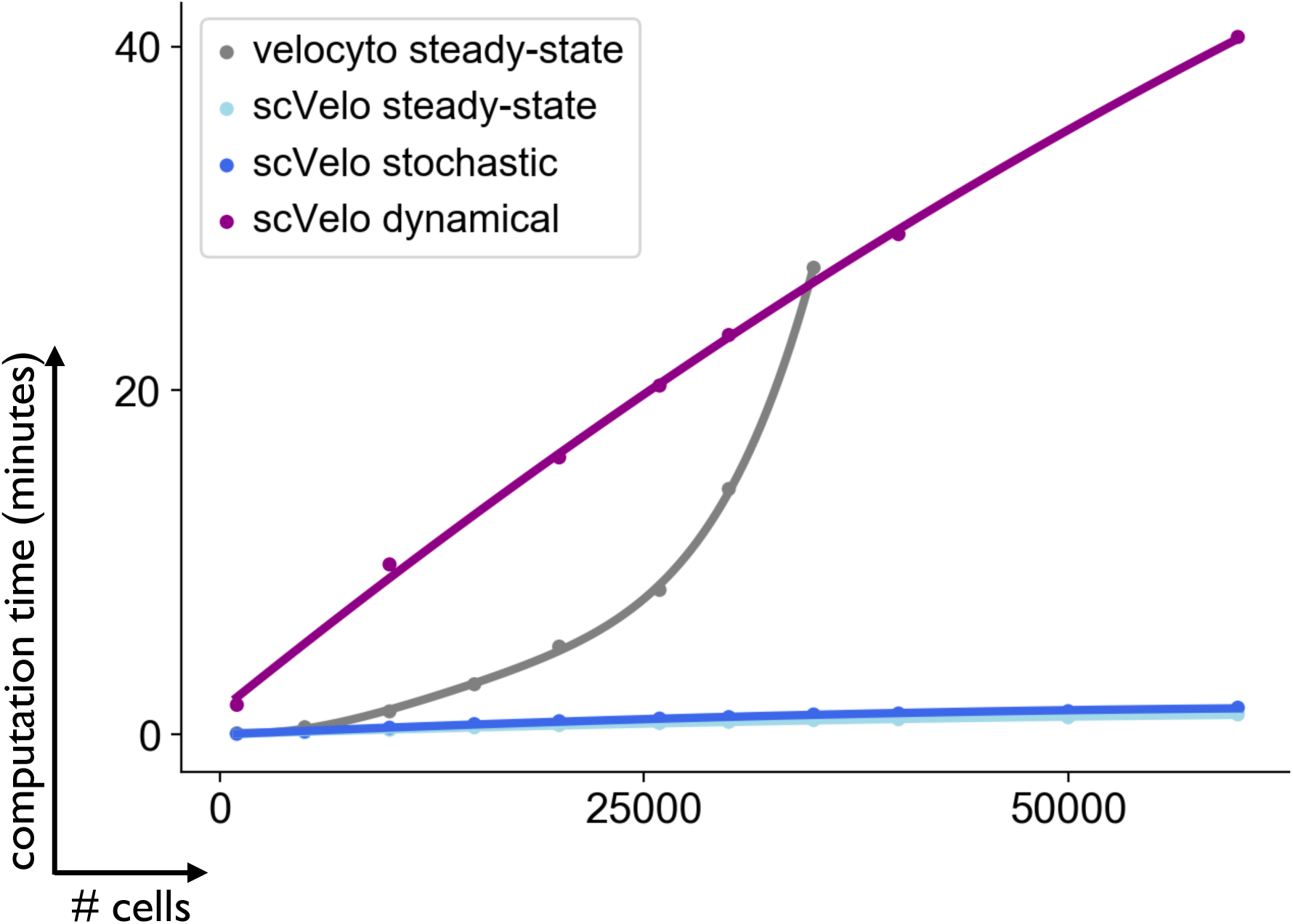
Runtime comparison. The dynamical, the stochastic as well as the steady-state model are available within scVelo as a robust and scalable implementation (https://scvelo.org). For comparison, we run the pipelines of velocyto’s steady-state model and all scVelo models on an Intel Core i7 CPU with 3.7GHz and 64GB RAM. We used 1k highly variable genes and varied the cell numbers from 1k to 60k cells. We further run the pipelines for 300k cells to test for scalability. The cells are sampled from the pancreas development dataset containing 25,919 transcriptome profiles. In less than a minute, scVelo manages to run the full velocity pipeline for the steady-state as well as stochastic model on 35k cells. The full splicing dynamics including kinetic rate parameters, latent time and velocities, is inferred in a longer but practicable runtime of 20 minutes. As it scales linearly with the number of cells and genes, its runtime is exceeded by velocyto’s quadratic runtime on large cell numbers of 35k and higher. For large cell numbers, also memory efficiency becomes a critical aspect. Velocyto cannot process more than 40k cells as it runs out of memory, while scVelo scales to more than 300k cells.

